# A recurrent network architecture explains tectal activity dynamics and experience-dependent behaviour

**DOI:** 10.1101/2022.03.30.486335

**Authors:** Asaph Zylbertal, Isaac H. Bianco

## Abstract

The ongoing activity of neuronal populations represents an internal brain state that influences how sensory information is processed to control behaviour. Conversely, external sensory inputs perturb network dynamics, resulting in lasting effects that persist beyond the duration of the stimulus. However, the relationship between these dynamics and circuit architecture and their impact on sensory processing, cognition and behaviour are poorly understood. By combining cellular-resolution calcium imaging with mechanistic network modelling, we aimed to infer the spatial and temporal network interactions in the zebrafish optic tectum that shape its ongoing activity and state-dependent responses to visual input. We showed that a simple recurrent network architecture, wherein tectal dynamics are dominated by fast, short range, excitation countered by long-lasting, activity-dependent suppression, was sufficient to explain multiple facets of population activity including intermittent bursting, trial-to-trial sensory response variability and spatially-selective response adaptation. Moreover, these dynamics also predicted behavioural trends such as selective habituation of visually evoked prey-catching responses. Overall, we demonstrate that a mechanistic circuit model, built upon a uniform recurrent connectivity motif, can estimate the incidental state of a dynamic neural network and account for experience-dependent effects on sensory encoding and visually guided behaviour.

## Introduction

Neural and behavioural responses to sensory input are influenced both by stimulus properties as well as the internal state of the brain, including the ongoing activity of neural networks. Consequently, repeated presentation of identical sensory cues evokes variable activity and this variation is left unexplained by analyses restricted to response averaging, as is traditionally used for empirically estimating neuronal tuning properties. Increasingly, however, such traditional approaches are being replaced by treating trial-to-trial variability as a product of continuous interactions between a dynamical network and external input (Churchland et al., 2011; Morcos and Harvey, 2016; Urai et al., 2019). One consequence of such interactions is the modulation of evoked responses by recent network activity (Arieli et al., 1996; Petersen et al., 2003; He, 2013; Shimaoka et al., 2019). Conversely, brief external input may perturb ongoing dynamics by pushing the network to a new state (Chen and Gong, 2019; Franco and Yaksi, 2021; Fritsche et al., 2021). This could have long-lasting outcomes beyond the duration of the stimulus itself, a memory effect influencing the responses to subsequent inputs (Fiser et al., 2004; Abbott et al., 2009; Deneux and Grinvald, 2016). These findings blur the distinction between ongoing and evoked processes in the brain and reinforce the view that both are governed by shared underlying network interactions (Curto et al., 2009; Bolt et al., 2018; Uddin, 2020). It is not clear, however, how these dynamic properties emerge from underlying circuit connectivity, nor how they modulate sensory encoding, perception and sensorimotor transformations.

Insights into such connectivity motifs and their role in network dynamics come from examining ongoing activity persisting in the absence of changes in the sensory environment. Such activity is commonplace in sensorimotor brain regions, including multiple cortical areas (Kenet et al., 2003; Yuste et al., 2005), and the optic tectum (Romano et al., 2015). Ongoing activity is typically not random but rather shows a characteristic correlation structure between individual neurons and across time (Tsodyks et al., 1999; Kenet et al., 2003; Smith et al., 2018). This structure is supported by recurrent connectivity, involving local and/or long-range feedback connections (Mohajerani et al., 2013; Zylbertal et al., 2017a; Smith et al., 2018). The spatiotemporal characteristics of ongoing activity are shaped by circuit properties such as connection strengths and vesicle release dynamics (Shu et al., 2006), as well as intrinsic cellular properties (e.g. ion channel dynamics, Zhang and Sillar, 2012; Zylbertal et al., 2015). A handful of stereotypical connectivity motifs encapsulating these properties are sufficient to explain ongoing dynamics (Haider et al., 2006; Galán, 2008; Betzel et al., 2019), and predict their consequences for sensory processing (Abbott et al., 2009; Curto et al., 2009; Rajan et al., 2010). However, to better understand the bidirectional interplay between ongoing activity and state-dependant sensory responses, it is necessary to test theoretical predictions arising from mechanistic models against experimental data within a common framework.

In this study, we analysed tectal activity data from larval zebrafish to estimate the principles of circuit organisation that underlie the interactions between ongoing and visually evoked activity and assessed the impact of such interactions on visuomotor behaviour. We first used light-sheet calcium imaging to observe ongoing activity in the complete tectal network and developed a stochastic, recurrent network model that reproduced the statistics of tectal activity. The model contains only two types of recurrent interactions that are applied equally to every neuron: fast local excitation and slow, long-range suppression. We then used our model to estimate the incidental state of every neuron in the network based on the recent history of spiking activity and found that this explains a portion of the trial-to-trial variability in responses to visual prey-like stimuli. The model also successfully predicted the reciprocal effect, where stimulus-evoked activity was followed by prolonged, spatially specific suppression of ongoing and visually evoked responses. These changes in tectal state were associated with modulation of spontaneous and visually-evoked prey-catching responses, suggesting a role for the recurrent tectal interactions in visuomotor transformations. In sum, a mechanistic network model, defined by a uniform recurrent connectivity rule, is able to explain how past activity influences the current state of the tectal network and thereby modulates visual representations and behaviour.

## Results

### The spatiotemporal structure of ongoing activity in the optic tectum

We reasoned that examining the spatial and temporal organisation of ongoing tectal activity was likely to be informative about the dominant connectivity motifs in the network. To measure this activity, we performed light-sheet calcium imaging of transgenic larvae expressing GCaMP6s (elavl3:H2B-GCaMP6s, 6 dpf, n=14), initially under constant environmental conditions (i.e. without presenting visual stimuli). We imaged a 375×410×75 μm volume encompassing the optic tectum (OT) and adjacent brain structures, at a rate of 5 volumes per second (Figure 1A-B). Following motion correction, each imaging plane was automatically segmented to individual neurons (Kawashima et al., 2016), and spikes were inferred by deconvolution of the extracted fluorescence signal (Friedrich et al., 2017). Imaged volumes were registered to a reference brain (Avants et al., 2009; Marquart et al., 2015) and somata in the stratum periventriculare (SPV), which comprise the majority of tectal neurons, were considered for further analysis (Figure 1B, 1.4·10^4^ ± 3·10^3^ tectal SPV cells per fish, mean ± SD).

**Figure 1:**
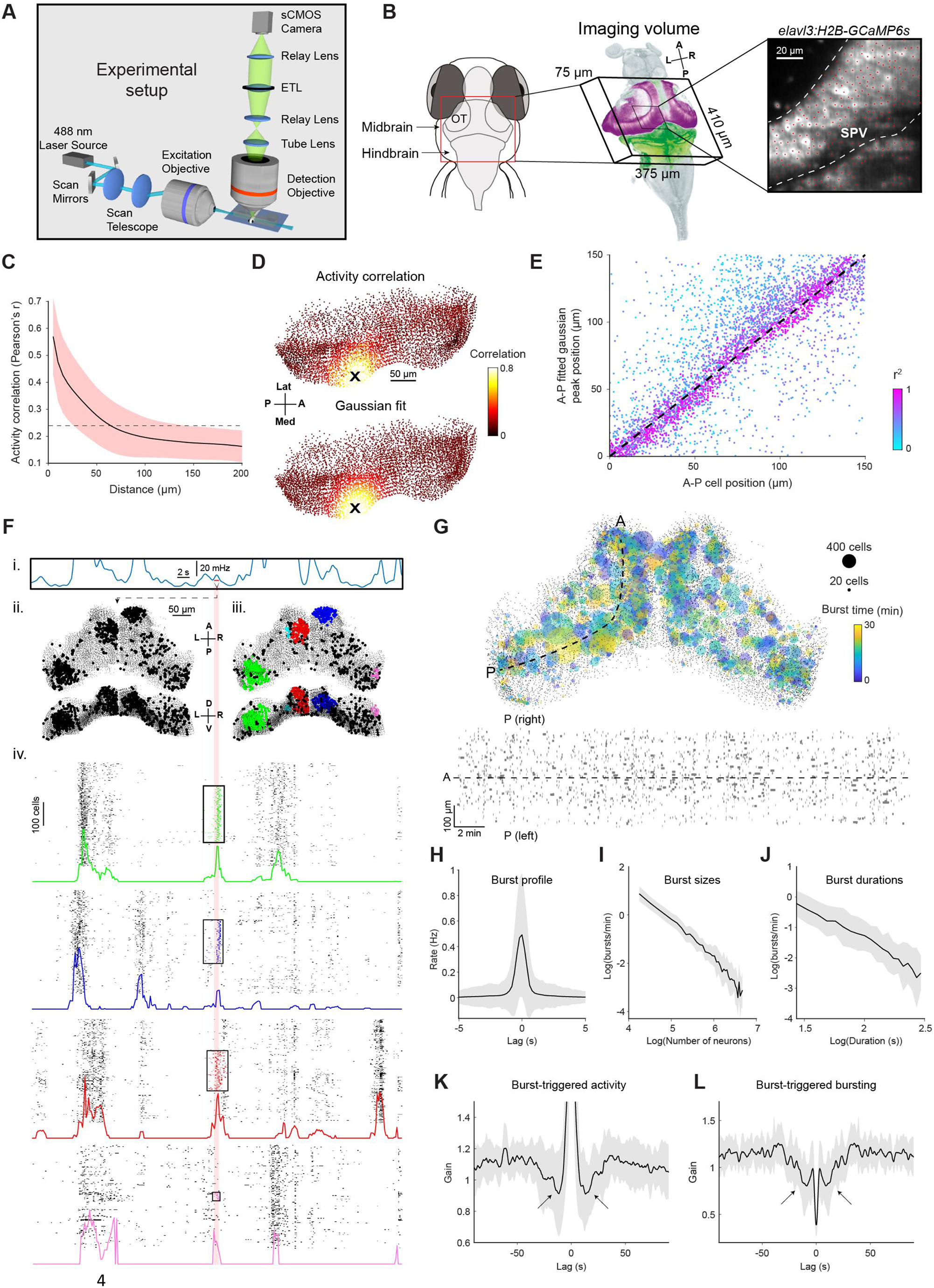
Tectal activity displays a uniform spatial correlation structure and localised bursting. (**A**) Light-sheet calcium imaging of tethered larval zebrafish (not to scale). (**B**) Left: Imaging field-of-view (green with tectal SPV mask in magenta) following registration to a reference brain (grey). Right: Section of a single imaging plane showing centroids of identified neurons (red). (**C**) Pairwise activity correlation as a function of Euclidean distance between cells in a single fish. Red shaded area denotes SD across seed cells (n=14597). Dotted line indicates mean correlation. (**D**) Map showing Pearson’s correlation between the activity of an example seed cell (X) and all other cells (top), and corresponding Gaussian fit (bottom). (**E**) Location of fitted correlation peaks vs location of seed cells in the same fish as D. Colour indicates r^2^ of Gaussian fits. (**F**) Example illustrating burst detection procedure. (i) Population activity peaks detected from mean spiking activity across all tectal neurons. (ii) Active cells identified within a 1 s window centred on an activity peak (red line). (iii) Active cells clustered according to their spatial density. (iv) Burst initiation and termination times determined for each cluster using Poisson-filtered activity of constituent neurons. Raster plots show the clustered cells participating in a burst as well as 200 neighbouring cells. Coloured traces show Poisson-filtered activity and rectangles denote the extent of each burst in time and space. (**G**) Top: Locations of all bursts detected during a single 30 min imaging session. Colours indicate burst times and spot sizes indicate number of participating neurons. Dashed line indicates the curved axis used for inferring A-P coordinates in (D-E). Bottom: Locations of burst centroids along A-P axis versus burst times. Rectangle widths indicate duration of each burst. (**H**) Mean activity of burst-participating neurons, centred on the peak of the Poisson-filtered activity. (**I-J**) Histogram of burst sizes (number of participating neurons, I) and durations (J). The rates of both quantities vary according to a power law, resulting in straight lines in a log-log plot. (**K**) Post-burst activity suppression: mean spiking of burst-participating neurons triggered on burst time. Data is normalised by mean activity of each neuron triggered on randomly sampled times. Arrows indicate post-burst activity suppression. (**L**) Probability of participation in a burst, triggered on burst time. Data is normalized for each fish by random circular permutation of burst times. Arrows indicate post-burst suppression of burst participation. Plots in H-L show mean values with grey shaded areas indicating SD across fish (n=14). ETL, electrically focus-tunable lens; OT, optic tectum; A, anterior; P, posterior; L, left; R, right; D, dorsal; V, ventral; Med, medial; Lat, lateral. See also Figure 1 – figure supplement 1.

The spatial pattern of activity correlations was compatible with recurrent excitatory interactions between tectal cells. To show this, we analysed the Pearson correlation between the ongoing activity of each neuron (‘seed’), and the activity of all other neurons during the entire experimental session. Correlation coefficients smoothly and monotonically decreased as a function of the Euclidean distance between neurons (Figure 1C and Figure 1 – figure supplement 1A-C). We confirmed this finding using 2-photon imaging, suggesting it is not a consequence of light scattering in the light-sheet microscope but rather a feature of tectal activity (Figure 1 – figure supplement 1C). We projected neuronal coordinates into two dimensions (anterior-posterior and medial-lateral tectal axes, see Materials and Methods), producing a 2D correlation map for each seed neuron (Figure 1D). A 2D gaussian was fitted to each map, excluding the seed neuron itself, yielding good fits for most cells (r^2^=0.42 ± 0.17, mean ± SD, Figure 1D and Figure 1 – figure supplement 1A-C). The peaks of the fitted gaussians showed close correspondence to the locations of seed neurons (Figure 1E and Figure 1 – figure supplement 1A-C). These data are compatible with a recurrent network in which excitatory interactions smoothly decay with distance from each cell.

A salient aspect of tectal activity is the occurrence of spontaneous bursts, during which spatially compact groups of neurons briefly fire in synchrony (Romano et al., 2015; Avitan et al., 2017). Such bursting might arise from recurrent excitatory interactions and so we sought to characterise the statistics of bursting to develop a model of tectal spatiotemporal interactions. To detect bursts, we evaluated the spatial distribution of cells active during one-second windows centred on population activity peaks (Figure 1F, i-ii). For each window, we identified the cells participating in a burst using density-based DBSCAN clustering (Ester et al., 1996, Figure 1F, iii and Figure 1 – figure supplement 1D-E, Materials and Methods). The temporal extent of each burst was then defined as the period of non-zero Poisson-filtered activity of participating neurons (Figure 1F, iv). Overall, 54424 bursts were identified in 14 fish. On average, each activity peak was associated with 1.5 ± 2.0 localised bursts (mean ± SD), bursts occurred at a rate of 46 ± 11 per minute and were uniformly distributed in space and time (Figure 1G). Bursts were brisk, phasic events, with the majority of spiking occurring over a timecourse of around one second (Figure 1H).

The statistics of bursting suggested a second, inhibitory network interaction shapes tectal activity. As has been previously reported, the sizes and durations of bursts were highly variable (95 ± 183 neurons per burst, 2.5 ± 3 s duration, mean ± SD), and distributed according to a power law (Figure 1I-J), suggestive of a critical phenomenon with ‘avalanche’ dynamics (Ponce-Alvarez et al., 2018). A universal feature in such phenomena is a self-limiting element responsible for both event termination and suppression of subsequent events. Therefore, we analysed how participating in a burst influences subsequent spiking activity (Figure 1K) and burst participation probability (Figure 1L) for individual neurons. Cells were less active and less likely to take part in another burst for tens of seconds. These observations are compatible with activity-dependent inhibition that has a self-limiting effect on bursting as well as a lasting effect extending beyond burst termination.

In sum, ongoing tectal activity is compatible with two general spatiotemporal features that might shape network dynamics. First, activity correlation is suggestive of a uniform connectivity motif in which excitatory recurrent connectivity smoothly decays with distance. Second, the dynamics of bursting suggests prolonged activity-dependent suppression.

### A spiking network model with a uniform connectivity rule reproduces tectal bursting

Next, we used computational modelling to explore the possibility that tectal bursting emerges from the recurrent connectivity motifs suggested by our analysis of ongoing activity. Theoretical network analysis has proposed that recurrent excitatory connectivity, balanced by long-range inhibition, produces stable, self-sustaining and spatially-defined ‘blobs’ of activity (Amari, 1977). We reasoned that by making the inhibition in such a model depend on a slow time-integration of activity, transient bursts would emerge instead of stable blobs and would be followed by prolonged activity suppression.

To test this idea, we used a probabilistic spiking network simulation based on a linear-nonlinear-Poisson (LNP) formalism. This framework has been successfully used to relate activity in neuronal populations to past activity and sensory input (Pillow et al., 2005, 2008; Truccolo et al., 2005), and its non-deterministic aspect fits the apparent stochastic nature of tectal bursting. In LNP models, the instantaneous firing rate of each neuron (λ) is determined by an exponentiated ‘linear drive’, computed as the sum of filtered inputs from all neurons in the network plus optional external inputs (Figure 2A, Pillow et al., 2008). Spikes are emitted by simulating a stochastic Poisson process according to the firing rate. Our model is thus defined by a set of seven parameters, describing the spatial and temporal aspects of excitatory (‘E’) and inhibitory (‘I’) interactions in the network. The connection weights between all pairs of neurons decay with Euclidian distance according to Gaussian functions with variances *σ*^(*E*)^, *σ*^(*I*)^, scaled by fixed gains *g*^(*E*)^, *g*^(*I*)^. Both interactions exponentially decay in time according to time constants *τ*^(*E*)^, *τ*^(*I*)^. A fixed bias term *μ* is summed with all other inputs to determine the total linear drive (see Materials and Methods). The locations of neurons in the simulation were taken from a single fish registered to the reference brain. Notably, given that the same connectivity rule applies to every tectal cell, the network is fully described by the set of seven parameters and the locations of the neurons.

**Figure 2:**
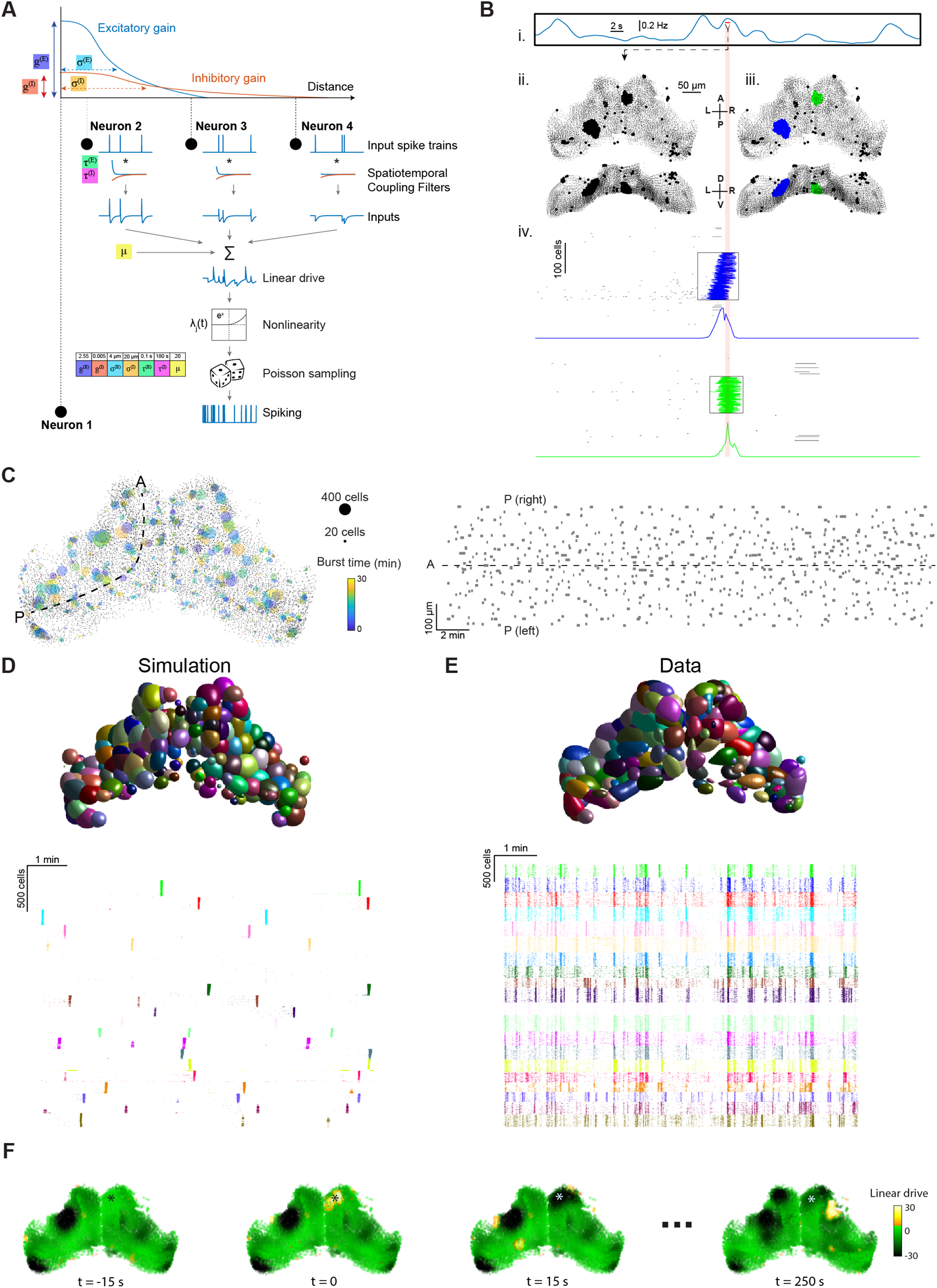
Stochastic spiking network model reproduces tectal bursting. (**A**) LNP Model architecture: connection weights for excitatory (E) and inhibitory (I) interactions are determined by Gaussian functions of the intercellular Euclidian distance, with unique gain (*g*^(*E*)^, *g*^(*I*)^) and spatial standard deviation (*σ*^(*E*)^, *σ*^(*I*)^) for each type of interaction. Presynaptic spikes are filtered in time with time constants (*τ*^(*E*)^, *τ*^(*I*)^) and summed along with a bias (*μ*) to produce the linear drive. Exponentiating this linear drive sets the mean of an inhomogeneous Poisson process from which spikes are randomly emitted (Materials and Methods). Model parameters used for all panels in this figure are shown inset. (**B**) Example of burst detection in the simulation results (c.f. Figure 1F). (**C**) Left: Locations of bursts detected during a 30 min simulation. Colours indicate burst times and spot sizes indicate number of participating neurons. Right: Burst locations vs. burst time. Rectangle widths indicate the temporal extent of each burst. (**D-E**) Neuronal assemblies detected using PCA-promax algorithm (Materials and Methods), for simulation results (D) and experimental data (E), shown on a spatial map (top) and a raster plot for representative assemblies (bottom). (**F**) Time-course of linear drive to model cells around the time of a spontaneous burst (t=0). Star indicates burst centroid.

In accordance with our hypothesis, running this network simulation resulted in spatially localised tectal bursting (Video 1). Bursting was a robust emergent property that was observed across a broad range of parameter values (see below). Bursts could be identified by analysing the simulation spiking output using the same method as for experimental data (described above) and were uniformly distributed in space and time with a range of sizes and durations (Figure 2B-C).

**Video 1: Spiking network model reproduces localised bursting**

Example simulation. Spiking neurons are depicted as yellow dots, neurons participating in a detected burst are marked by fading circles. Quiescent periods have been omitted. A, anterior; P, posterior; L, left; R, right; D, dorsal; V, ventral.

In previous studies, localised bursting in the zebrafish OT has been attributed to sub-networks with enhanced connectivity, termed neuronal assemblies (Romano et al., 2015; Avitan et al., 2017; Marachlian et al., 2018; Mölter et al., 2018; Diana et al., 2019). A PCA-promax algorithm was previously used to identify such assemblies based on the tendency of neurons to be co-active (Romano et al., 2015). We applied this method to our simulated tectal network and despite the fact that, by design, there were no subgroups of cells with preferential connectivity, PCA-promax yielded multiple spatially localised `assemblies` (Figure 2D), as was also the case for recorded data (Figure 2E). This demonstrates that repeated, synchronous bursting of tectal neurons neither implies nor requires preferentially connected sub-networks but can instead emerge in a network with uniform distance-based recurrent connectivity.

Our simulated tectal network enabled us to examine ‘sub-threshold’ activity of simulated neurons (specifically, their linear drive), in addition to spiking output. The population linear drive before, during and after bursts (Figure 2F, Video 2) reveals, as expected, that bursts tend to occur in regions with higher than average linear drive (bright green). Moreover, bursts leave behind a region of low linear drive (black), supressing the emergence of a burst in the same region for a long period of time.

In sum, our network model reveals that a uniform recurrent connectivity motif is sufficient to explain ongoing tectal bursting.

**Video 2: Simulated linear drive**

Time-course of linear drive to model cells in a single simulation run.

### Model optimization reproduces the statistics of tectal bursting

We next evaluated the capacity of the model to produce bursting with biologically relevant statistics. We found that bursting occurred robustly under a range of parameter values, while statistics such as average burst size and duration varied across parameter space. For example, starting from the set of parameters used in the previous section (‘baseline model’) and systematically varying the inhibitory space constant (*σ*^(*I*)^), inhibitory time constant (*τ*^(*I*)^) and global bias (*μ*) produced a wide range of mean burst frequencies (0.5 – 165 bursts per minute) and sizes (55 – 230 neurons per burst, Figure 3A).

**Figure 3:**
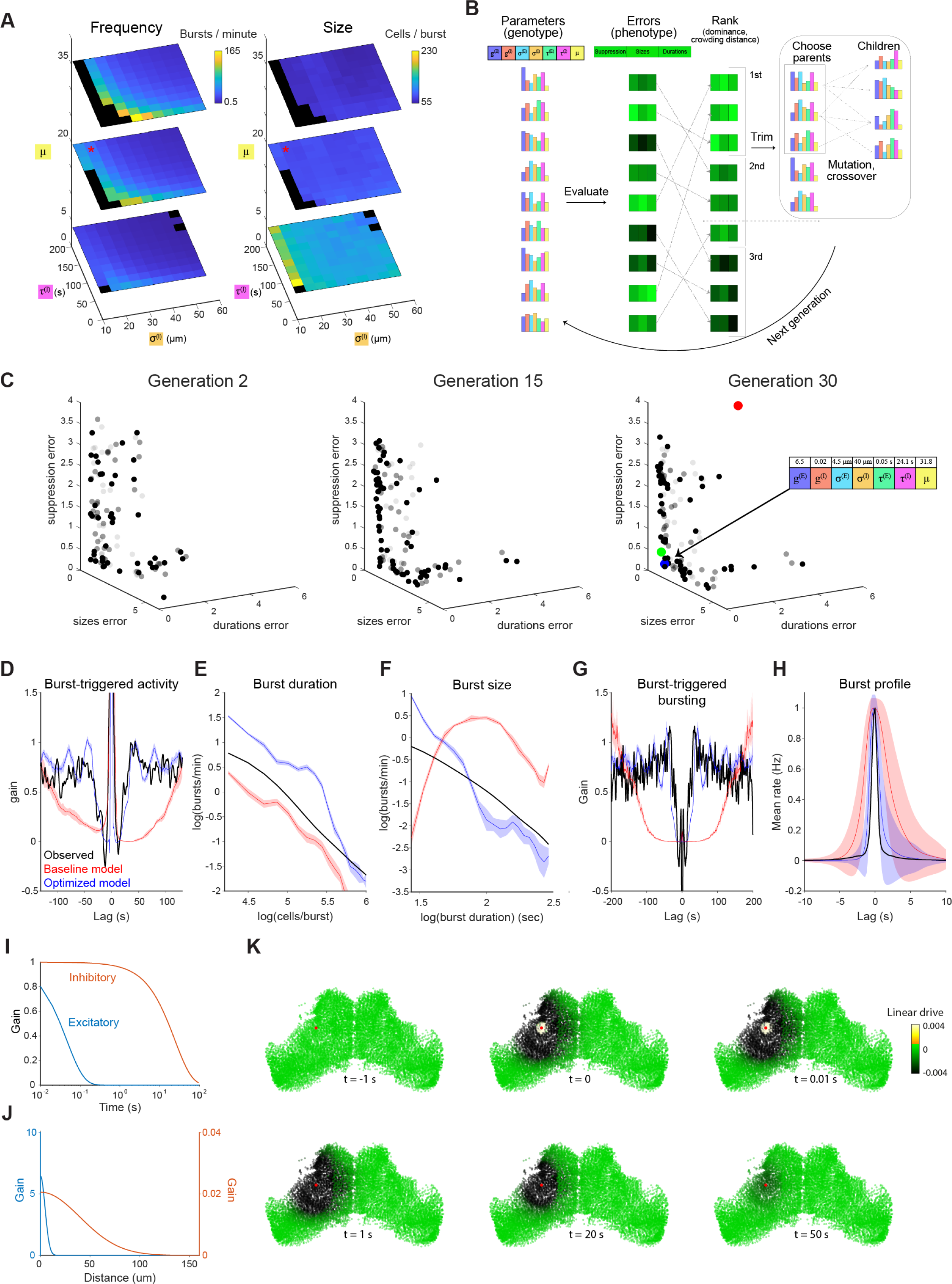
Optimization of model parameters using EMOO. Burst rate (left) and number of participating neurons (right) as a function of inhibitory space constant (s^(I)^), inhibitory time constant (τ^(I)^) and bias (μ). Red star indicates the parameters used in baseline model (as per Figure 2), and black regions indicate parameter combinations that failed to produce bursting. (**B**) Schematic of EMOO (evolutionary multi-objective optimization). In each generation, a population of models are evaluated for their error scores on the three objectives and top-ranking models are selected. New models (‘children’) are produced by crossover and mutation from ‘parents’ and the extended population (selected top-ranking models and children) forms the next generation (see Materials and Methods). (**C**) Model populations of the 2^nd^ (left), 15^th^ (middle) and 30^th^ (right) EMOO generations, shown in the z-scored error space. Bold spots indicate the non-dominated set (first rank set), faint and fainter spots indicate the second and third rank sets, respectively. Red spot: baseline model; green spot: elbow solution used to initialise pattern search; blue spot: final optimized model (parameters indicated on the right). (**D-F**) Model results evaluated against optimization objectives: post-burst activity suppression (D), burst sizes distribution (E) and burst durations distribution (F). (**G-H**) Model predictions vs experimental data for two additional features: Burst-triggered burst participation (G, see Figure 1L) and burst temporal profile (H, see Figure 1H). (**I-J**) Temporal (I) and spatial (J) profiles of the intercellular interactions in the optimised model. (**K**) Timeseries illustrating the influence of a single spike (in cell marked with red spot) on the linear drive of surrounding cells in the optimised model network. Plots show mean and shaded areas indicate SD for n=10 simulation runs.

Next, we sought to fit the model parameters to match observed tectal bursting statistics. We used evolutionary multi-objective optimization (EMOO, Deb, 2001, Figure 3B and Materials and Methods) to tune the parameters according to three target objectives, namely the distribution of burst sizes (Figure 1I) and durations (Figure 1J), and the timecourse of burst-triggered activity suppression (Figure 1K). With each successive generation, EMOO produces a population of models that better approximates the Pareto optimal set of solutions (Pareto front), i.e. the set of models where no individual model performs better than any other in all three objectives (Figure 3C). Following thirty EMOO generations, we chose the solution at the elbow of the estimated Pareto front (the model closest to the origin in the z-scored loss space, Figure 3C, green), and used it as a starting point for a local search to fine-tune the parameters.

The resulting optimized model was successful in reproducing the bursting statistics of the zebrafish tectum (Figure 3C, blue). This was true for both the features used as optimization objectives (Figure 3D-F), as well as two that were not directly fit, namely burst-triggered burst participation probability (Figure 3G) and the burst temporal profile (Figure 3H).

The fitted parameter values of the optimized model support the idea that bursting in OT arises from distinct spatiotemporal patterns of excitatory versus inhibitory network interactions. Excitatory connections are strong, short range and decay rapidly, whereas inhibitory connections extend over longer distances and decay slowly and thus integrate activity for long periods of time (Figure 3I-J). As a result, a single spike in a given neuron strongly excites its immediate neighbours for a brief moment (< 1 s), while slightly reducing linear drive across a large part of the tectal hemisphere for tens of seconds (Figure 3K).

Because our optimized network model could reproduce spontaneous activity with biologically realistic statistics, we next explored the extent to which it could account for bidirectional interactions between incidental network state and stimulus-evoked activity.

### Incidental network state influences visually evoked responses in OT

The dynamical state of neuronal populations is likely to contribute to variability in neural representations of sensory stimuli. To explore this, we applied our network model to recorded tectal activity data in order to estimate the incidental state of the biological network immediately prior to individual visual stimulus presentations and thereby predict the influence of ongoing activity upon stimulus-evoked responses.

We presented zebrafish with small moving spots, which evoke robust tectal activity (Figure 4A, Niell and Smith, 2005; Bianco and Engert, 2015). Spots swept horizontally across the visual field at two elevations, one slightly below and one above the horizon and to mitigate interactions between consecutive stimuli we initially used a long inter-stimulus interval (ISI, 5 min). The responses of tectal neurons displayed retinotopic mapping and direction selectivity (Figure 4B, Stuermer, 1988; Niell and Smith, 2005; Hunter et al., 2013).

**Figure 4:**
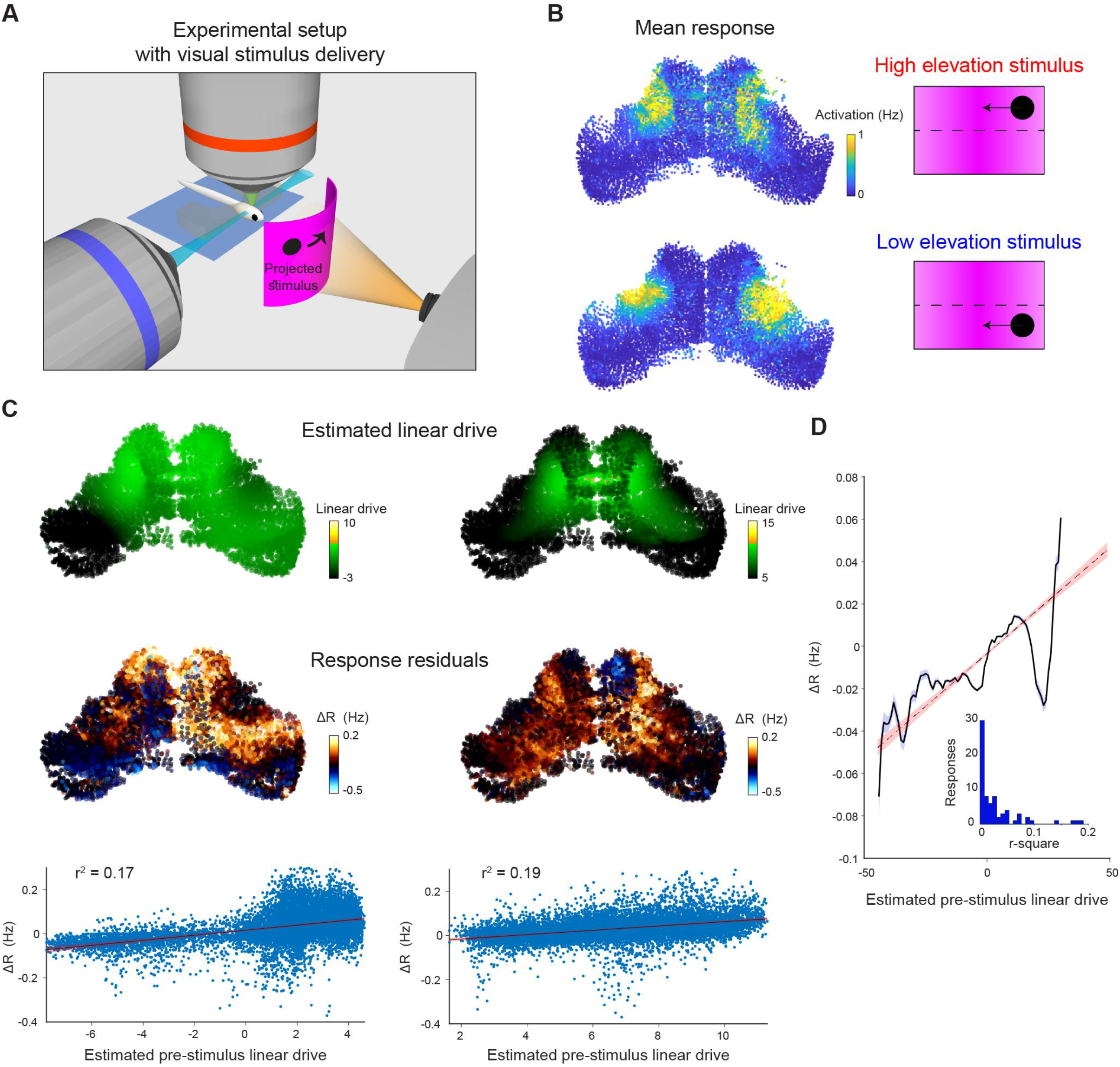
Incidental network state influences visually evoked responses. **(A)** Visual stimuli comprising prey-like moving spots were presented to larvae during light-sheet imaging. (**B**) Average firing rate of tectal cells during presentation of leftwards moving spots at high (top) and low (bottom) elevation. (**C**) Two examples of stimulus presentations showing model-estimated pre-stimulus linear drive (top), recorded response residuals (middle), and residuals versus linear drive (bottom). (**D**) Response residuals as a function of linear drive for 72 stimulus responses in 6 fish. Grey shaded area indicates SEM. Dashed line with red shading indicates linear fit with confidence intervals for α = 10^−10^. Inset: Distribution of r^2^ values for the 72 trials.

Using this data, we examined the relationship between response variability and model-estimated network state. First, we computed for each neuron the response residual for each stimulus presentation as the difference between the response for that specific stimulus presentation and the trial-averaged response to the stimulus.

Next, we used our optimised model to estimate the incidental state of OT neurons (their ‘linear drive’) just prior to each stimulus presentation, based on the recent history of ongoing activity in the network. Spiking activity, inferred from calcium imaging data during the minute prior to stimulus onset (specifically, −60 to −5 s before stimulus onset) was spatiotemporally filtered and summed for each cell using the optimised model parameters (Materials and Methods).

We found that estimated linear drive was positively correlated with response residuals (Figure 4C). This was true both for individual stimulus presentations and when we pooled data across 72 responses from 6 fish (Figure 4D).

These results suggest that the spatiotemporal interactions implemented within our network model can predict incidental network state and thereby account for a fraction of the trial-to-trial variability of visually evoked tectal activity. When the recent activity history is such that neurons are predicted to be in a state of higher excitability, this tends to result in a greater-than-average response to external sensory input.

### Ongoing activity is supressed following visually evoked activity

Next, we explored the reciprocal interaction, namely if stimulus-evoked activity perturbs ongoing network dynamics and if such interactions are predicted by our network model.

To do this, we first expanded our model to incorporate external (sensory) input such that model neurons were activated according to the visual receptive fields of their corresponding tectal neurons. We estimated receptive fields by fitting Gaussian functions to the measured activity of each cell as a function of stimulus angle (Figure 5A, Materials and Methods). Receptive fields collectively covered visual space, with an average width (2.5σ) of 38° ± 24° (mean ± SD, Figure 5B), comparable to previous reports (Niell and Smith, 2005). Because visual space is retinotopically mapped within OT, neurons responsive to high and low elevation stimuli occupied distinct anatomical locations (Figure 5C, Materials and Methods).

**Figure 5:**
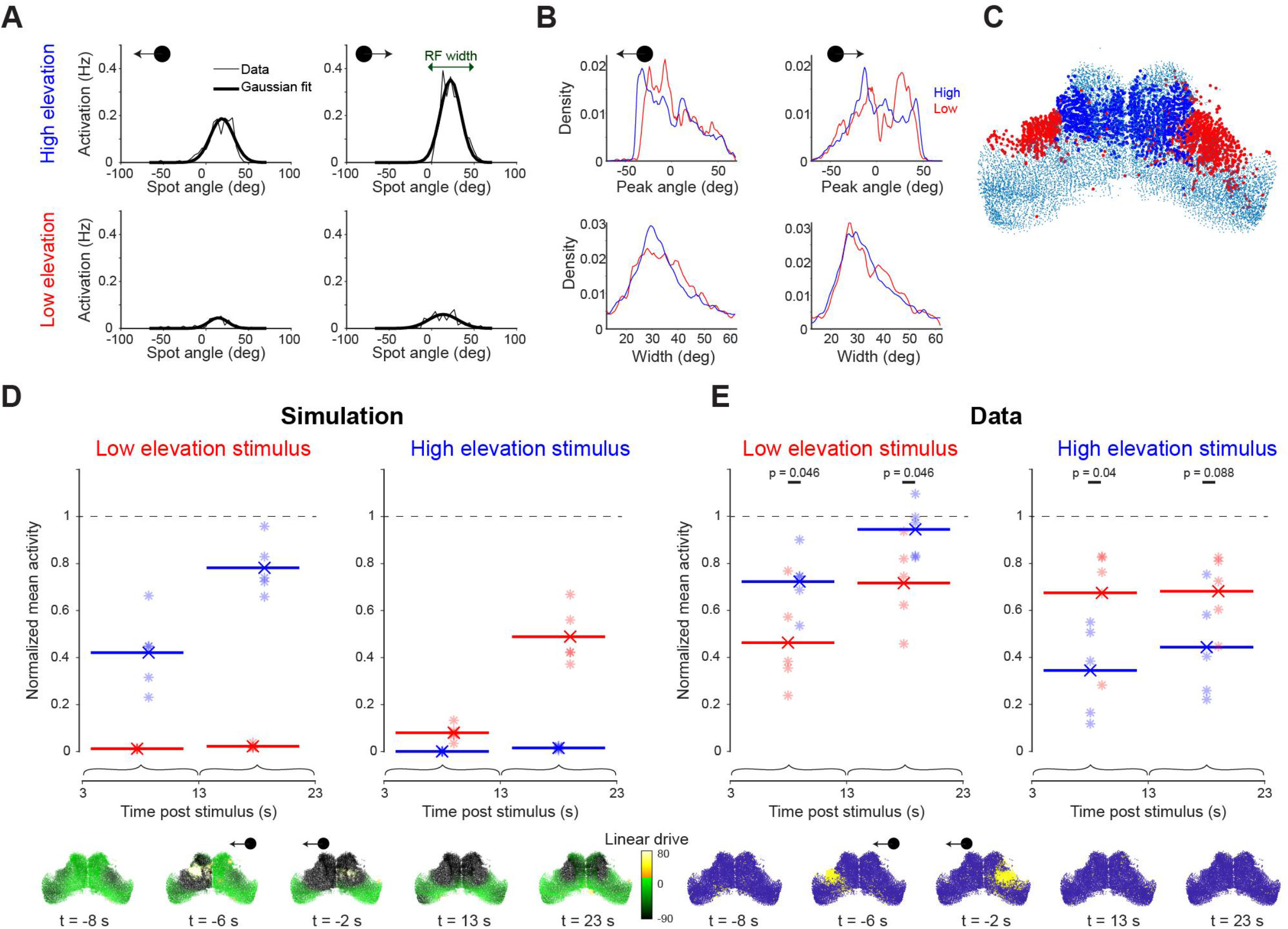
Visual stimulation is followed by long lasting, spatially selective suppression of tectal activity. **(A)** Visually evoked activity of an example neuron showing mean spike counts (thin line) and Gaussian fit (thick line) for moving spots at high (top) and low (bottom) elevation and moving leftward (left) or rightward (right). This neuron shows modest direction selectivity. (**B**) Distribution of receptive field centres (top) and widths (bottom) for cells with good Gaussian fits (r^2^ > 0.8). (**C**) Locations of cells responsive to high (blue) and low (red) elevation stimuli. (**D-E**) Top: Post-stimulus activity for simulated (D) and recorded (E) neurons. Activity is shown for cells responsive to high (blue) and low (red) elevation and integrated across two time windows following the cessation of the visual stimulus. Each point indicates a single simulation run or experimental session, lines indicate average values. Values are normalized by the mean ongoing activity. Bottom: linear drive (D) and recorded activity (E) for single example trials at the indicated times from stimulus offset.

Running the network simulation with external sensory input predicted a substantial, long-term reduction in tectal activity for tens of seconds following visual stimulation (Figure 5D). This prediction is concordant with the long inhibition time constant in the model and, as a result of the spatially weighted recurrent connectivity in the model, was biased to recently stimulated regions of OT. Thus, cells responsive to the recently presented stimulus were predicted to show a greater suppression of ongoing activity in the period following stimulus offset as compared to neurons responsive to the other elevation (Figure 5D). This spatial-selective suppression of activity was explained by a lasting state of low linear drive extending beyond the duration of the sensory stimulus (Figure 5D, Video 3).

**Video 3: Simulated linear drive during and after visual stimulus**

Time-course of linear drive to model cells in a simulation of an evoked response to a moving spot stimulus (black circle).

To test this prediction, we repeated the same analysis with calcium imaging data from five fish. As predicted by the model simulation, neurons responsive to the recently presented stimulus showed a marked reduction of ongoing activity for tens of seconds, whereas neurons responsive to the other elevation showed a lesser suppression of post-stimulus activity (Figure 5E).

In sum, our network model can account for bidirectional interactions between ongoing and evoked tectal activity. The incidental state of the network modulates visually evoked responses, while these responses, in turn, exert a long-lasting suppressive effect upon ongoing activity.

### Long lasting activity dependent inhibition can explain adaptation of visual responses

These observations lead to the prediction that visual stimuli will have a lasting effect on the state of the tectal network that may influence subsequent sensory responses. Thus, we next used network modelling and calcium imaging to explore how the recent history of sensory stimulation impacts visually evoked activity and behaviour.

Based on the timescales and spatial ranges apparent in tectal dynamics, we designed an experimental paradigm incorporating a mixture of temporal and spatial relationships between visual stimuli. During 30 minute ‘stimulation blocks’, small moving spots were presented at two different elevations (out of a possible three). One stimulus was presented at high frequency (‘common’, 30 s inter-stimulus interval, ISI) and the second at low frequency (‘deviant’, 5 min ISI). Each deviant stimulus was followed by a 90 s break. Stimulation blocks were separated by 30 minute ‘rest blocks’, during which no stimuli were presented (Figure 6A).

**Figure 6:**
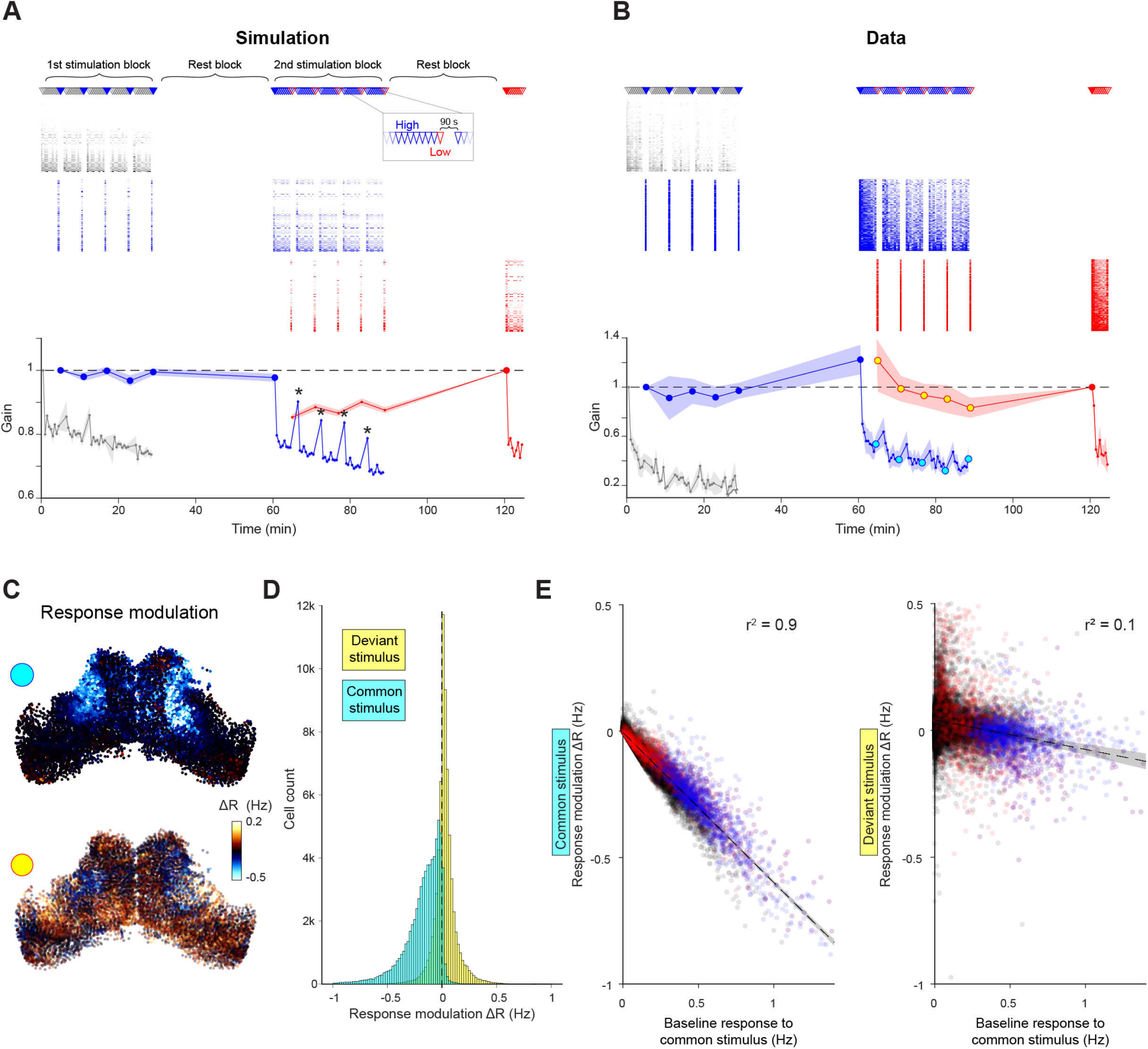
Selective adaptation to visual stimuli. (**A-B**) Simulated (A) and recorded (B) responses of cells tuned to very low (grey), high (blue) and low (red) elevation stimuli. Top: Stimulation protocol. Middle: Raster of responses of individual cells. Bottom: Mean response across cells tuned to each elevation, normalised by first response. Large symbols indicate epochs used to establish the `baseline` response for analysis in (C-E), stars indicate partial response recovery following 90 s breaks. Shaded areas indicate SEM for n=5 fish or simulation runs. **(C)** Top: Response modulation for the high elevation stimulus. For every OT cell we compare the baseline response (large blue circles in (B)) to the 2^nd^ stimulation block (cyan circles in B) where the stimulus is presented as a common stimulus. Bottom: Response modulation for the low elevation stimulus, comparing baseline responses (large red circle in (B)) to the 2^nd^ stimulation block (yellow circles in (B)) where it is presented as a deviant stimulus. (**D**) Distribution of single cell response modulation for the common (cyan) and deviant stimulus (yellow), (n = 67750 cells from 5 fish). (**E**) Single cell response modulation for common (left) and deviant stimulus (right) as a function of the baseline response to the common stimulus. Blue and red spots indicate cells tuned to the common and deviant stimuli, respectively. Shaded areas indicate linear regression confidence intervals for α = 10^−10^. See also Figure 6 – figure supplement 1.

We first evaluated the model prediction for this stimulus protocol by simulating it five times. To take into account adaptation of retinal inputs, we imaged the tectal neuropil in *isl3:GAL4;UAS:SyGC6s* fish expressing GCaMP6s at the synaptic terminals of retinal ganglion cells (RGCs), and modulated the input gain for the simulated high frequency stimulus accordingly (Figure 6 – figure supplement 1 and Materials and Methods). The model predicted an adaptation of tectal visual responses to the common stimulus, but not to the deviant stimulus (Figure 6A). For example, during the 2^nd^ stimulation block, high and low elevation stimuli were presented as common and deviant stimuli respectively (Figure 6A, blue and red). Adaptation reduced the mean response of cells tuned to the high elevation stimulus to about 70% of their initial amplitude followed by a partial recovery during the 90 s breaks (Figure 6A, stars). This is due to a combination of gradual input (RGC) adaptation and additional activity dependent suppression intrinsic to the tectum (Figure 6 – figure supplement 1C-D). Conversely, the response to the low elevation stimulus remained approximately constant.

To test these predictions against experimental observations, we used the same stimulation protocol with five fish (Figure 6B). In agreement with the model prediction, high-frequency stimulus presentation caused a spatially specific response adaptation to the common stimulus. The adaptation pattern matches the predicted combined effect of RGC adaptation and intrinsic tectal dynamics (Figure 6A), rather than either one alone (Figure 6 – figure supplement 1C-D).

To examine how this response modulation varies across cells, we compared the average response of individual cells during the 2^nd^ stimulation block to their baseline response (i.e. their average response to low-frequency stimulation, large circles in Figure 6B). The response to the common (high elevation) stimulus was attenuated in most cells (Figure 6C-D, cyan), and attenuation magnitude was linearly correlated with the amplitude of the baseline response (Figure 6E). By contrast, response modulation to the deviant (low elevation) stimulus was less consistent (Figure 6C-D, yellow) although some attenuation was observed in cells that were strongly activated by the common stimulus (Figure 6E). These results show that the common stimulus drives a subset of tectal cells to a supressed state, which modulates their subsequent responses to either stimulus.

To conclude, OT displays experience-dependent response modulation as predicted by our network model. This effect allows for integration of past stimuli and suppression of subsequent responses to similar stimuli.

### Tectal activity dynamics explain modulation of prey-catching behaviour

Lastly, we examined how network dynamics relate to visually guided prey-catching behaviour, which is mediated by OT (Gahtan et al., 2005; Muto et al., 2013; Bianco and Engert, 2015; Helmbrecht et al., 2018). Because hunting routines of larval zebrafish invariably commence with eye convergence, we used the occurrence of convergent saccades (CS) to identify prey-catching behavioural responses (Figure 7A inset, Bianco et al., 2011; Trivedi and Bollmann, 2013; Bianco and Engert, 2015).

**Figure 7:**
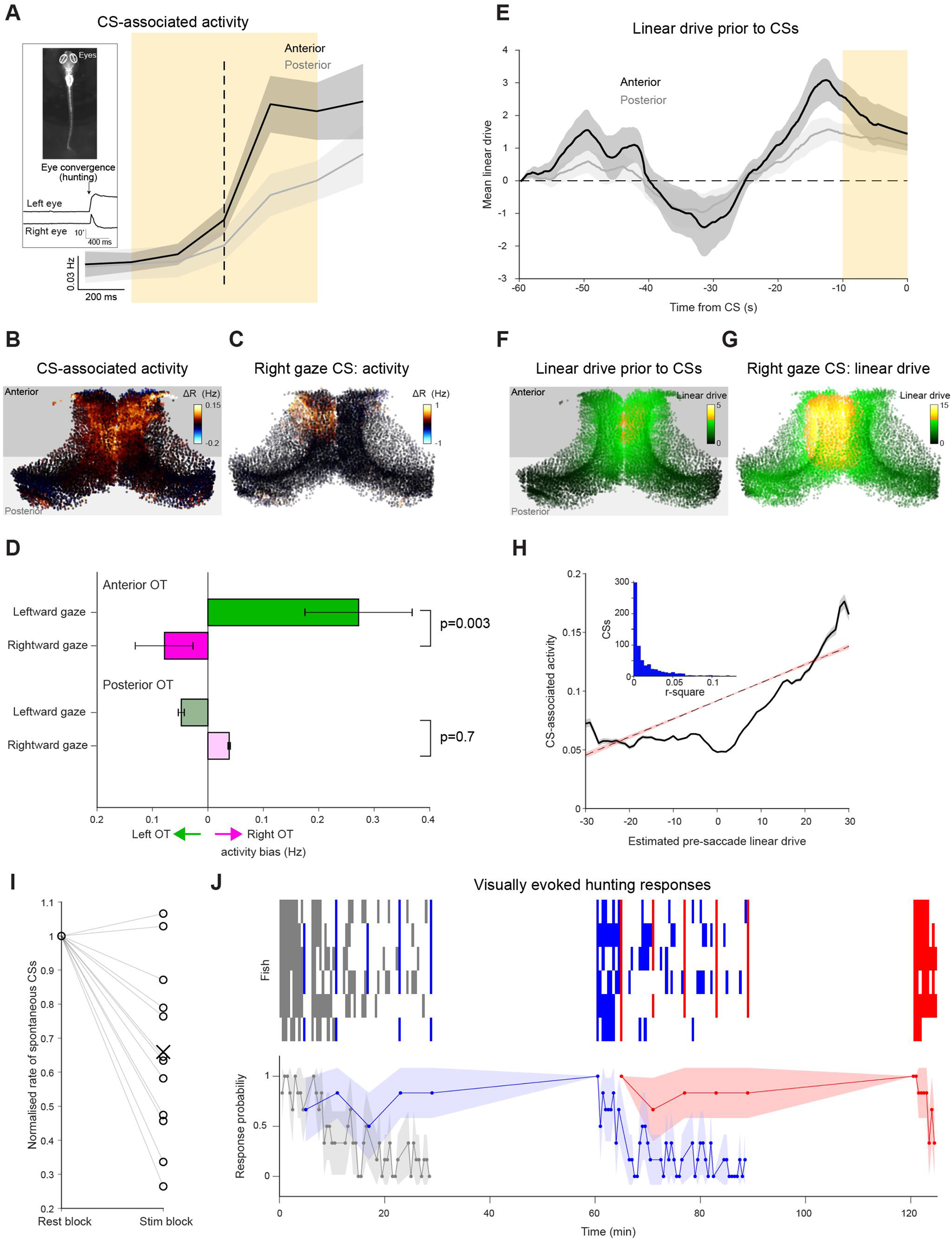
Prey-catching behaviour is modulated by tectal network state and stimulus history. (**A**) Activity of neurons in the anterior (black) and posterior (grey) OT, triggered on spontaneous CSs. Mean with shaded areas indicating SEM (n=11 fish), dashed line indicates CS time. Inset: Eye tracking and an example of a detected convergent saccade. (**B**) Mean baseline-subtracted activity of OT cells during 1-s windows centred on spontaneous CSs (orange rectangle in (A), n=66 CSs in a single fish). (**C**) Baseline-subtracted activity for a single spontaneous CS with rightward post-saccade gaze angle. (**D**) Comparisons of activity between OT hemispheres for lateralised CSs with leftward (green) and rightward (magenta) post-saccade gaze angle. Error bars indicate SEM (n=408 CSs from 11 fish). (**E**) Linear drive in the anterior (black) and posterior (grey) OT, prior to CSs. Mean with shaded area indicating SEM (n=11 fish). Dashed line indicates baseline linear drive. (**F**) Linear drive of OT cells during the 10 s prior to spontaneous CSs (orange rectangle in (E), in the same fish depicted in (B), n=66 CSs). (**G**) Linear drive prior to the spontaneous CS shown in (C). (**H**) CS-associated activity of individual cells as a function of estimated pre-CS linear drive (680 spontaneous CSs in 11 fish). Dashed line indicates linear fit to all single cell responses. Grey shaded area indicates SEM. Inset: Distribution of r^2^ values for the 680 CSs. (**I**) Rate of spontaneous CSs during rest blocks (more than 120 seconds since the last visual stimulus presentation) vs visual stimulation blocks, normalised by rest block rate. Each line indicates one fish. (**J**) Prey-catching responses evoked by very low (grey) high (blue) and low (red) elevation stimuli, presented according to the protocol described in Figure 6. Top raster show responses from each of 6 animals (rows) and lower trace shows mean response probability with 90% confidence interval.

We first analysed “spontaneous” CSs, which occasionally occur in the absence of visual stimuli (Bianco & Engert, 2015), to assess the link between ongoing tectal activity and behavioural output. By examining activity in 1 s time windows centred on CSs (‘CS-associated activity’, Figure 7A, orange shaded area), we found that spontaneous CSs are associated with an increase in tectal activity, particularly in the anterior tectum (Figure 7A-B, anterior tectum p=0.002, n=11 fish, Wilcoxon signed rank test, anterior vs posterior activity p=0.02, n=11 fish, Wilcoxon signed rank test). Moreover, CS-associated activity was lateralised in accordance with the direction of behaviour. We showed this by comparing activity between the tectal hemispheres for spontaneous CSs that directed gaze to the left or right (>10° post-saccade gaze angle). In anterior (but not posterior) OT, CS-associated activity was significantly biased towards the hemisphere contralateral to post-saccadic gaze (Figure 7C-D, p=0.003 and 0.7 for anterior and posterior OT, n=408 CSs, two-way ANOVA). These findings are in agreement with previous reports linking anterior OT activity with prey-catching behaviour directed to the contralateral side (Bianco & Engert, 2015; Fajardo et al., 2013; Helmbrecht et al., 2018).

These observations are compatible with intrinsic tectal dynamics contributing to the generation of spontaneous prey-catching responses. Alternatively, CS-associated activity could be a consequence of the eye movement (visual or proprioceptive sensory feedback), or represent a motor efference copy. Therefore, to investigate if tectal activity is likely to cause spontaneous behaviour, we used our network model to estimate the linear drive state of tectal neurons immediately prior to spontaneous CSs based on the recent activity history of the network (same method as above, Figure 4). This analysis showed that linear drive in anterior OT was significantly elevated prior to CSs (Figure 7E-F, pre-CS vs baseline p=0.004, n=11 fish, one-tailed t-test), and this difference was larger compared to posterior OT (p=0.04, n=11 fish, one-tailed paired t-test). At a single cell level, pre-CS linear drive was positively correlated with CS-associated activity, both for individual CSs (Figure 7G) as well as for data pooled over all spontaneous CSs (Figure 7H, compare Figure 4D). These results are compatible with ongoing dynamics in the tectal network contributing to the generation of spontaneous prey-catching behaviour.

Because we have shown that the recent history of visual stimuli has lasting effects upon tectal activity, we next asked how past experience of visual inputs modulates spontaneous and visually-evoked prey-catching behaviour. To do this, we used the mixed-ISI stimulation protocol (Figure 6), while tracking behaviour.

When we analysed spontaneous behaviour, we found that the rate of spontaneous CSs in the two minutes that followed any visual stimulus was reduced by approximately one third compared to rest blocks (Figure 7I, n=13 fish, p=0.002, Wilcoxon signed rank test). This suppression of behaviour is concordant with the lasting suppression of linear drive and ongoing tectal activity described above. Moreover, the probability of visually-evoked prey-catching was modulated by stimulus history in a qualitatively similar way to visually evoked tectal activity. Specifically, response rates declined for common, but not deviant stimuli (Figure 7J, compare Figure 6B).

Taken together, these observations suggest that the state of the tectal network, reflecting the integration of previous inputs and ongoing activity, modulates both spontaneous and visually evoked prey-catching behaviour.

## Discussion

In this study, we combined functional imaging and network modelling to explore the bidirectional interactions between ongoing and evoked neural activity and infer the dominant aspects of network architecture that underlie these dynamics. Based on our observations of ongoing activity, we inferred a recurrent connectivity motif, incorporating fast local excitation and long-lasting activity-dependent suppression, which was able to reproduce the occurrence and statistics of tectal bursting. By applying the same spatiotemporal interactions to recorded activity, we showed that they predict a network state that forms the background against which visual input is processed, explaining a fraction of the variability in visually-evoked responses. The model also predicted how visual stimuli perturb tectal dynamics to produce experience-dependent suppression of both ongoing and visually-evoked activity. Finally, comparisons of predicted network state to prey-catching responses suggest these dynamics modulate tectally-mediated behaviour. Our modelling and experimental findings suggest that a uniform recurrent connectivity motif can account for multiple aspects of ongoing dynamics and state-dependent visuomotor responses.

### Bursting activity and the tectal assemblies hypothesis

Recent studies have suggested that the bursting activity observed in OT represents recruitment of sub-networks with enhanced connectivity, or neuronal ‘assemblies’ (Romano et al., 2015; Avitan et al., 2017; Marachlian et al., 2018; Mölter et al., 2018; Diana et al., 2019). At the core of this model is the idea that there must exist nonuniformities in connectivity in which cells forming an assembly are preferentially interconnected and operate as a local attractor network. On the basis of assumptions such as these, various methods have been developed to detect assemblies from patterns of ongoing tectal activity (Mölter et al., 2018).

However, this study calls these assumptions into question. We show that bursting in OT is reproduced in a network model where there are no such subnetworks but rather the same connectivity rule (in which recurrent connection strengths smoothly decay according to Gaussian functions) is implemented for every cell. Although such homogeneous connectivity is almost certainly an oversimplification, this simple model nonetheless reproduced multiple aspects of tectal activity including the spontaneous bursting that has previously been interpreted as the signature of assemblies (Figure 2D).

Moreover, the assembly model does not account for observations such as the continuous distribution of bursts sizes even in a limited tectal region (Figure 1G) and the lack of discernible ‘cliques’ in the pair-wise activity correlation structure (Figure 1D-E). While it is very likely that anisotropies in connectivity will underlie more complex aspects of tectal physiology, we argue that inferring heterogeneities in connectivity on the basis of spatially localised bursting is not robust and should be revisited.

### The biological implementation of functional connectivity motifs

The recurrent connectivity motifs in our network model predict that the activity of a given OT neuron transiently and strongly increases the excitability of its immediate neighbours, while causing a sustained suppression of a broader population of neurons in its vicinity (Figure 3K). How might these functional connections be implemented in the tectal circuit?

The short time scale we estimated for excitatory interactions (*τ*^(*E*)^ = 0.05 *s*) is compatible with direct, recurrent synaptic connectivity in OT. Such connections have been demonstrated by intracellular recording of delayed excitatory post-synaptic potentials in tadpole tectum following optic tract stimulation (Pratt et al., 2008). Delayed glutamatergic potentials were similarly recorded in slices from rat superior colliculus and are thought to mediate pre-saccadic bursting (Saito and Isa, 2003).

By contrast, the inhibitory interactions in our model caused activity-dependent suppression lasting tens of seconds (*τ*^(*I*)^ = 24.1 *s*), too long to be accounted for by direct ionotropic synaptic inhibition. This interaction might be mediated by pre-synaptic depression, which can persist for tens of seconds in the avian OT (Luksch et al., 2004). Alternatively, slow activity suppression may be mediated by activity-dependent changes in intracellular properties such as ionic concentration or conductances (‘intrinsic plasticity’), potentially lasting for several minutes (Desai et al., 1999; Karmarkar and Buonomano, 2006; Gasselin et al., 2015; Zylbertal et al., 2017b).

Long term suppression might also be mediated by tonic activity in inhibitory populations. There is currently no evidence to suggest that tectal interneurons are the source of such suppression, since their visually-evoked activity appears to be limited to transient responses (Preuss et al., 2014; Dunn et al., 2016). Alternatively, brain regions interconnected with OT, such as GABAergic populations in the vicinity of the isthmic hindbrain (Gebhardt et al., 2019; Henriques et al., 2019), might provide long lasting feedback inhibition.

### Tectal dynamics and visuomotor transformation underlying hunting behaviour

Our results indicate that the state of the tectal network is predictive of prey-catching response probability, in agreement with the role of OT in generating premotor signals for this visuomotor behaviour (Herrero et al., 1998; Gahtan et al., 2005; Bianco and Engert, 2015; Antinucci et al., 2019). What adaptive purpose might the network interactions we describe play during natural behaviour?

OT, and its mammalian homolog the superior colliculus, is thought to be involved in detection and selection of salient inputs and generation of rapid behavioural responses (Boehnke and Munoz, 2008; Dutta and Gutfreund, 2014). The experience-dependent activity suppression we observe is compatible with this role, as it filters out expected inputs which are less likely to be associated with immediate danger or prey-catching opportunity. Similarly, zebrafish OT exhibits response adaptation to looming stimuli that is thought to play a role in the habituation of escape behaviour (Marquez-Legorreta et al., 2019). Comparable stimulus-selective adaptation and behavioural habituation has been described for auditory stimuli in barn owl OT (Reches and Gutfreund, 2008; Netser et al., 2011), and for overhead looming stimuli in the mouse superior colliculus (Lee et al., 2020). Notably, adaptation in the barn owl is selective for multiple auditory stimulus features such as interaural time and level differences, as well as frequency and amplitude. In the simple model we propose, the strength of interactions between neurons depends solely on Euclidean distance and therefore the spatial proximity of their receptive fields. An interesting avenue for future study will be to examine whether recurrent interactions weighted by similarity of feature tuning can account for such feature-selective adaptation.

## Materials and Methods

### Animals

Zebrafish (*Danio rerio*) larvae were reared on a 14/10 hr light/dark cycle at 28.5°C. For all experiments, we used zebrafish larvae homozygous for the *mitfaw2* skin-pigmentation mutation (Lister et al., 1999). For Ca2+ imaging experiments, we used larvae homozygous for *Tg(elavI3:H2B-GCaMP6s)*^*jf5Tg*^ (Vladimirov et al., 2014, ZFIN ID: ZDB-ALT-141023–2). For imaging RGC axonal projections, larvae were double transgenic for *Tg(Isl2b:Gal4)*^*zc60Tg*^ (Fredj et al., 2010, ZFIN ID: ZDB-ALT-101130-1) and *Tg(UAS:SyGCaMP6s)*^*a155Tg*^ (Dunn et al., 2016 ZFIN ID: ZDB-ALT-160406-1). All larvae were fed *Paramecia* from 4 dpf onward. Animal handling and experimental procedures were approved by the UCL Animal Welfare Ethical Review Body and the UK Home Office under the Animal (Scientific Procedures) Act 1986, under Home Office project licence 70/8162 awarded to Isaac Bianco.

### Light-sheet functional calcium imaging and behavioural tracking

For calcium imaging we used a custom-built digitally scanned light-sheet microscope. The excitation path included a 488 nm laser source (OBIS, Coherent, Santa Clara, California), a pair of galvanometer scan mirrors (Cambridge Technology, Bedford, Massachusetts) and objective (Plan 4X, 4x/0.1 NA, Olympus, Tokyo, Japan). A water-immersion detection objective (XLUMPlLFLN, 20x/1.0 NA, Olympus), a tube lens (f=200 mm), two relay lenses (f=100 mm) in a 4f configuration, and sCMOS camera (Orca Flash 4.0, Hamamatsu, Hamamatsu, Japan) were used in the orthogonal detection path. For remote focusing (Fahrbach et al., 2013), an electrically tunable lens (ETL, EL-16-40-TC-VIS-20D, Optotune, Dietikon, Switzerland) was installed between the relay lenses, conjugate to the back focal plane of the objective. Volumes (375 × 410 × 75 μm) comprising 19 imaging planes spaced 4 μm apart, were acquired at 5 volumes/s. Each plane received laser excitation for 1 ms (duty cycle 9%) resulting in average laser power at sample of 12.4 μW. To keep the observed population of neurons in each plane in focus throughout long imaging sessions, we implemented an automatic correction for slow drift in the Z direction. At the beginning of the experiment we acquired two reference stacks, centred on two of the imaging planes, by incrementally biasing the Z scanning mirror and ETL in steps of 1.5% of their scan amplitude. During the course of the experiment, Z drift was estimated every 30 seconds by comparing recent images to these reference stacks, finding the reference images with the maximal XY cross-correlation, and averaging the two drift estimates. Every five minutes the Z scan mirror and ETL signals were biased to offset any detected drift, according to the average of the ten most recent Z drift estimates.

For functional imaging, larval zebrafish were mounted in a custom 3D printed chamber (SLS Nylon 12, 3DPRINTUK, London, United Kingdom) in 3% low-melting point agarose (Sigma-Aldrich, St. Louis, Missouri) at 5 dpf and allowed to recover overnight before functional imaging at 6 dpf. Visual stimuli were back-projected (ML750ST, Optoma, New Taipei City, Taiwan) onto a curved screen forming the wall of the imaging chamber in front of the animal, at a viewing distance of ∼10 mm. A coloured filter (Follies Pink No. 344, Roscolux, Stamford, Connecticut) was placed in front of the projector to block green light from the collection optics. Visual stimuli were designed in Matlab (MathWorks, Natik, Massachusetts) using Psychophysics toolbox (Brainard, 1997). Stimuli comprised 10° dark spots on a bright magenta background, moving at 20°/s either left→right or right→left across ∼110° of frontal visual space. Two or three elevation angles were used, calibrated for each fish during preliminary imaging by finding elevations separated by at least 15° that produce robust observable tectal activation (typically a very low elevation stimulus ∼25° below the horizon, a low elevation stimulus ∼10° below the horizon, and a high elevation stimulus ∼5° above the horizon).

Eye movements were tracked during imaging experiments at 50 Hz under 850 nm illumination using a sub-stage GS3-U3-41C6NIR-C camera (Point Grey, Richmond, Canada). The angle of each eye was inferred online using a convolutional neural network (three 5×5 convolutional layers with 1, 1 and 4 channels, each followed by stride-2 max pooling layer, and a single fully connected layer), pre-trained by annotating images from multiple fish covering a wide range of eye positions. Eye movements were categorized as a convergent saccade if both eyes made nasally directed saccades within 150 ms of one another. Microscope control, stimulus presentation and behaviour tracking were implemented using LabVIEW (National Instruments, Austin, Texas) and Matlab.

### 2-photon imaging

Functional 2-photon calcium imaging was carried out as described in Antinucci et al., 2019 to record tectal activity in a single imaging plane in the absence of visual stimulus presentation for 30 min. Prior to imaging initiation, the larva was allowed to adapt to the imaging rig for 15 min.

### Calcium imaging analysis

All calcium imaging data analysis was performed using Matlab scripts. Volume motion correction was performed by 3D translation-based registration using the Matlab function ‘imregtform’, with gradient descent optimizer and mutual information as the image similarity metric. A registration template was generated as the time-average of the first 10 volumes and then iteratively updated following each block of 10 newly registered volumes from the first 500 frames. This template was then used to register all remaining volumes. For elavl3:H2B-GCaMP6s experiments, 2D regions of interest (ROIs) corresponding to cell nuclei were computed from each template imaging plane using the cell detection code provided by (Kawashima et al., 2016). For RGC axonal arbor imaging, two ROIs encompassing the tectal neuropil were manually defined for each imaging plane. The time-varying raw fluorescence signal *F*_*raw*_*(t)* for each ROI was extracted by computing the mean value of all pixels within the ROI mask at each time-point. A slowly varying baseline fluorescence *F*_*0*_*(t)* was estimated by taking the 10^th^ percentile of a sliding 20-volume window, and was used to calculate the proportional change in fluorescence:

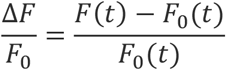

These values were subsequently zero-centred by subtracting the mean for each ROI, and ROIs with a slow drift in their baseline fluorescence (for which the standard deviation of mean-normalised *F*_*0*_*(t)* was greater than 0.45) were discarded.

To estimate spike trains, the zero-centred 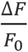 time series for each cell was deconvolved using a thresholded OASIS (Friedrich et al., 2017) with a first-order autoregressive model (AR(1)) and an automatically-estimated transient decay time constant for each ROI.

To standardize the 3D coordinates of detected cell nuclei, template volumes were registered onto the *Tg(elavl3:H2B-RFP)* reference brain in the ZBB brain atlas (Marquart et al., 2017) using the ANTs toolbox version 2.1.0 (Avants et al., 2011) with affine and warp transformations. As an example, to register the 3D image volume in ‘fish1_01.nrrd’ to the reference brain ‘ref.nrrd’, the following parameters were used:

~~~
antsRegistration -d 3 -float 1 -o [fish1_, fish1_Warped.nii.gz] -n
BSpline -r [ref.nrrd, fish1_01.nrrd, 1] -t Rigid[0.1] -m
GC[ref.nrrd, fish1_01.nrrd, 1, 32, Regular, 0.25] -c
[200×200×200×0,1e-8, 10] -f 12×8×4×2 -s 4×3×2×1 -t Affine[0.1] -m
GC[ref.nrrd, fish1_01.nrrd, 1, 32, Regular, 0.25] -c
[200×200×200×0,1e-8,10] -f 12×8×4×2 -s 4×3×2×1 -t SyN[0.1,6,0] -m
CC[ref.nrrd, fish1_01.nrrd, 1, 2] -c [200×200×200×200×10,1e-7,10]
-f 12×8×4×2×1 -s 4×3×2×1×0
~~~

Following registration, tectal ROIs were labelled using a manually created 3D mask (Supplementary file 1).

### Burst detection

We used the same procedure to detect localised bursts of tectal activity in both recorded and simulated data. First, we found local maxima in tectal population activity after averaging the inferred spike trains of all tectal cells and smoothing it with a sliding window 3 frames in width (600 ms). We next identified the population of cells that were active during a 1-second window centred on each peak, and used the DBSCAN density-based clustering algorithm (Ester et al., 1996) to define spatial clusters based on Euclidean distances between the centroids of active cells. We excluded peaks during episodes of strong bilateral activation (commonly associated with vigorous swims or struggles) where more than 10% of the total tectal neurons were active and less than 70% of the active cells were located in the same hemisphere (16% ± 5% population peaks excluded per fish). We set the clustering distance threshold ϵ to 15 μm (∼3 cell diameters) and the minimal number of cells in a neighbourhood, MinPts, to 12. MinPts was selected as the minimal number for which no clusters were detected after repeatedly performing circular permutations the time bases of all neurons, Figure 1 – figure supplement 1D-E). After detecting the clusters associated with an activity peak, each was analysed independently to determine the initiation and termination times of the bursting activity of its constituent cells. A sliding window, six imaging frames in width (1.2 s) was evaluated at single frame increments. At each position, spike counts were discarded if they exceeded the 60^th^ percentile of a Poisson cumulative distribution function with λ equal to the mean number of inferred spikes across all cells in the cluster. Finally, the mean number of remaining spikes was computed and used to produce a vector describing the average Poisson-filtered firing within the cluster. Its first and last non-zero values were defined as the burst initiation and termination times.

### Activity correlation analysis

To assess the correlation structure of ongoing tectal activity, we calculated the Pearson correlation of the spiking activity of each neuron (‘seed’) and the activity of all other tectal neurons. Spiking vectors were smoothed by filtering with a Gaussian window with σ = 1.4 frames (280 ms). To generate correlation maps, we first projected the spatial coordinates of tectal neurons onto two dimensions: anterior-posterior and medial-lateral tectal axes. This was done by manually tracing a curve along the tectal anterior-posterior axis and calculating the distance of each neuron to this curve (its medial-lateral location) and the location along the curve where it passes closest to each neuron (its anterior-posterior location). Next, the two-dimensional correlation map of each seed (excluding the seed itself) was fitted to a general bivariate normal distribution density function:

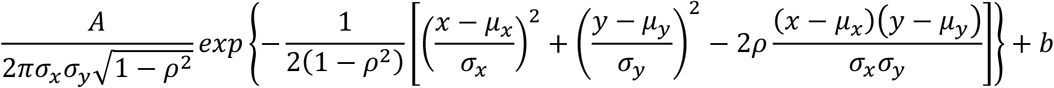

Where *A* is the gain, b is the bias, *μ*_*x*_and *σ*_*x*_ are the A-P axis mean and standard deviation, *μ*_*y*_and *σ*_*y*_ are the M-L axis mean and standard deviation, and *ρ* is the correlation coefficient.

A random location within 100 μm of the seed cell was used as an initial guess for the peak (*μ*_*x*_, *μ*_*y*_). The following bounds were used when fitting:

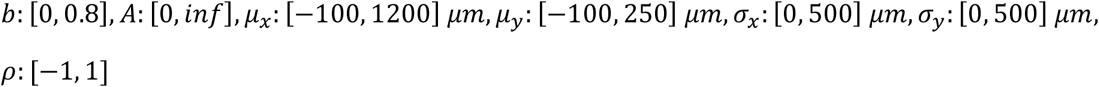

### LNP spiking network model

The network model was based on the LNP stochastic spiking network formalism (Pillow et al., 2005, 2008; Truccolo et al., 2005). Interactions between neurons (including feedback from a neuron onto itself) were modelled by temporal coupling filters and external (visual) input was considered instantaneous (see below).

The instantaneous firing rate of neuron j at time t, *λ*_*j*_(*t*), was derived by exponentiating its linear drive *φ*(*t*), or sum of inputs:

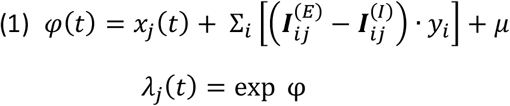

Where *x* is the external stimulus, 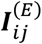 and 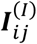 are excitatory and inhibitory coupling filters, *y*_*i*_ are spike train histories of each neuron at time *t*, and *μ* is a global baseline log-firing rate (DC input).

The excitatory and inhibitory coupling filters were each determined by three global parameters: gain (g), spatial influence standard deviation (σ) and temporal exponential decay time constant (τ).

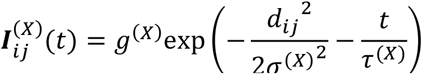

Where X denotes excitatory (E) or inhibitory (I) interactions and *d*_*ij*_ is the Euclidian distance between neuron *i* and neuron *j* in 3D standard coordinates following registration. If the two neurons reside in opposite tectal hemispheres, their coupling strengths were both scaled by 0.01.

The model simulation was implemented in Matlab (code available at https://github.com/azylbertal/TectalLNP), based on code provided by (Pillow et al., 2008). It uses a 50 ms timestep (resulting in 4:1 ratio with imaging rate) and comprises 14733 simulated neurons with locations inferred from the tectal SPV of one fish. To significantly reduce memory and computational costs, a much longer (5 s) time step was used to simulate the slow inhibitory inputs to all cells. The inhibitory input was then interpolated before being summed with the excitatory input to produce the total linear drive. Spikes were drawn at random based on an inhomogeneous Poisson process with expected rates *λ*_*j*_(*t*), which were updated according to the equations above whenever at least one neuron spiked. To compare the simulation results with imaging results, simulated spiking output was binned to 200 ms windows, equivalent to imaging volumes.

### Model fitting

We fit the seven model parameters (*g*^(*E*)^, *g*^(*I*)^, *σ*^(*E*)^, *σ*^(*I*)^, *τ*^(*E*)^, *τ*^(*I*)^, *μ*) to reproduce observed bursting characteristics using the Matlab implementation of controlled elitist evolutionary multi-objective optimization with a genotype-based crowding distance measure (the Matlab function ‘gamultiobj’, Deb, 2001). Specifically, we used three optimization objectives: a) the log-rates of bursts binned according to the logarithm of the number of participating neurons; b) the log-rates of bursts binned according to the logarithm of the burst duration; c) mean burst-triggered spiking activity (specifically, the average activity of each neuron triggered on the peaks of bursts it participated in and subsequently averaged across all neurons, smoothed by spline interpolation and scaled from zero to one according to its 5^th^ and 95^th^ percentiles).

In each generation of the algorithm, a population of (initially random) 200 models was evaluated in parallel using UCL’s Myriad computing cluster. Each simulation was run without external inputs for 306000 steps (the equivalent of two hours and 15 minutes of imaging), and the initial 18000 steps (15 minutes) were discarded to avoid initialisation transients. Localised bursts were detected based on the simulated spiking as they were detected for experimental data (see above). Since the inter-hemispheric interactions in the model are very weak (1% of the intra-hemispheric interactions), we were able to reduce computational cost during optimization by evaluating the models on one hemisphere. The loss value for each objective is the sum of squared errors between the simulation and experimental results. Parents for the next generation are selected by tournament: subsets of models are chosen by random (with replacement), and the best one from each set, i.e. the one with the highest ranking from the previous generation, is selected as a parent. 80% of the children are produced by intermediate crossover, where their parameters are chosen randomly from a uniform distribution bounded by the parameters of pairs of parents. The remaining 20% are produced by adaptive mutation, where they are randomly mutated with a direction that is maintained upon an increase in mean population fitness from the previous generation, or changed otherwise. The extended population (original population and children) is then evaluated and ranked based on Pareto front order, and within each front by genotype crowding distance (individuals with less close neighbours are ranked higher). Finally, the extended population is trimmed to retain the best 200 individuals for the next generation.

The evolution was stopped when the population scores stopped improving (∼30 generations), producing a set of models on the approximated Pareto front - the solutions for which no individual model performs better than any other in all objectives. We chose the solution at the ‘elbow’ of the resulting front (the one closest to the origin in the z-scored loss space) and used it as a starting point for a local pattern search for final parameter refinement (Figure 3).

### Linear drive estimation based on imaging data

We estimated the linear drive, or ‘excitability’ of each OT neuron prior to stimulus presentation (Figure 4) or spontaneous convergent saccade (Figure 7) by applying the spatiotemporal filters described by the LNP model to spiking history inferred from imaging data. For each event *i*, we calculated 60-s long vectors *φ*_*ij*_(*t*) representing the sum of inputs to each cell *j*, based on the activity of all other OT cells during the 60-5 s period prior to the event weighted by time and distance per the model parameters (Eq 1 above). Since the model temporal filters assign larger weights to recent spikes, we ignored the spikes detected during the final 5 seconds to account for the inherent uncertainty in inferring spikes from calcium imaging data. To account for cell-specific biases, such as spatial edge effects, we corrected *φ*_*ij*_(*t*) by subtracting cell-specific ‘baseline’ mean linear drive vectors 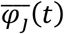, estimated by repeating the calculation for 200 randomly-sampled time points. The reported linear drive point-estimate was calculated by averaging the final 10 seconds of the baseline-corrected vectors.

### Receptive field mapping and visual stimulus simulation

The simulated external input to each model neuron, *x*_*j*_(*t*), was estimated from the mean recorded activity of the corresponding tectal neuron as a function of stimulus azimuth (*λ*_*j*_(*α*)). To estimate receptive fields, the following function was fitted to this data:

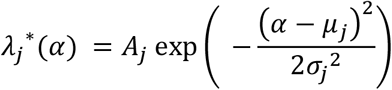

producing values for gain (*A*_*j*_), peak azimuth (*μ*_*j*_) and width (*σ*_*j*_) for a specific neuron, stimulus elevation and motion direction (Figure 5A-B). To reduce the impact of noise, neurons with very low peak response or very narrow (*σ*_*j*_<3°) receptive fields were excluded from receiving external input in the model.

Finally, to infer external input drive from measured spiking output, we scaled the gain and receptive field widths (to account for apparent expansion caused by recurrent connectivity) to obtain:

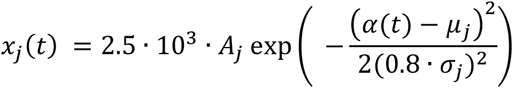

Our results are robust to substantial variations in the choice of these scaling factors.

### Identification of responsive neurons

We identified tectal neurons as responsive to one of the stimulus elevations (Figure 5-6) if their mean response firing rate exceeded the 90^th^ percentile over the responses of all neurons and was at least 5 times higher than their baseline firing rate.

### Statistical tests

p-values were obtained from two-tailed t-tests unless otherwise noted. For the two-way ANOVA used to model lateral activity bias (Figure 7D), post-saccadic gaze direction (rightward or leftward) and fish identity were used as independent factors, without interaction terms. The reported p-value is associated with the gaze direction.

## Data Availability

Neural activity and behavioural data have been deposited to Dryad, under doi:10.5061/dryad.q573n5tm5. Source data files have been provided for all figures. Simulation source code is available at https://github.com/azylbertal/TectalLNP.

## Acknowledgments

The authors thank members of the Bianco lab, James Fitzgerald, Eyal Wigderson and Avraham M Libster for helpful discussions and critical feedback on the manuscript and UCL Fish Facility staff for fish care and husbandry. A.Z. was supported by a BBSRC Discovery Fellowship (BB/S010564/1). I.H.B. was supported by a Wellcome Trust and Royal Society Sir Henry Dale Fellowship (101195/Z/13/Z) and a Wellcome Trust Senior Research Fellowship (220273/Z/20/Z).

---

**Figure 1 – figure supplement 1:**
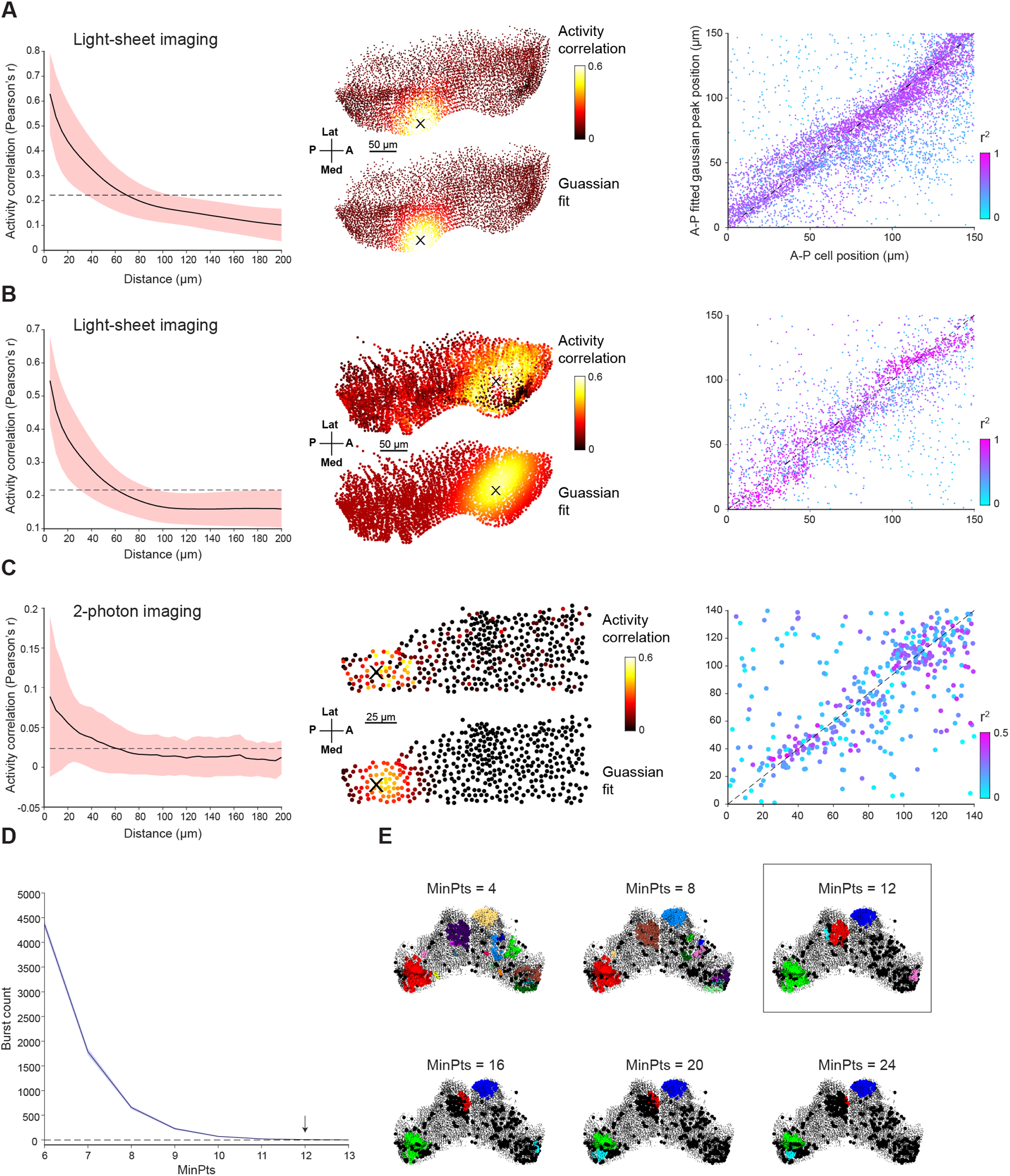
Ongoing activity and localised bursting in the optic tectum. (**A-B**) Left: Pairwise activity correlation as a function of Euclidean distance between cells in two additional example fish (red shaded area denotes SD across cells, n=18580 and 10948). Middle: Example correlation maps and Gaussian fits for the same fish. Right: Positions of fitted Gaussian peaks vs positions of seed cells. Spot colours indicate r^2^ values. (**C**) Same analysis as A-B but for 2-photon imaging of single plane in OT of a third fish. (**D**) To select the MinPts parameter for DBSCAN, we performed a shuffle analysis in which the time base for each neuron was circularly permuted. Plot shows the mean number of detected bursts across 10 permutations as a function of the MinPts parameter. Shaded area indicates SD, arrow indicates the conservative value we chose, where zero bursts were detected in shuffled data. (**E**) Bursts detected for one tectal activity peak for different values of MinPts. Rectangle indicates chosen value. A, anterior; P, posterior; Med, medial; Lat, lateral

**Figure 6 – figure supplement 1:**
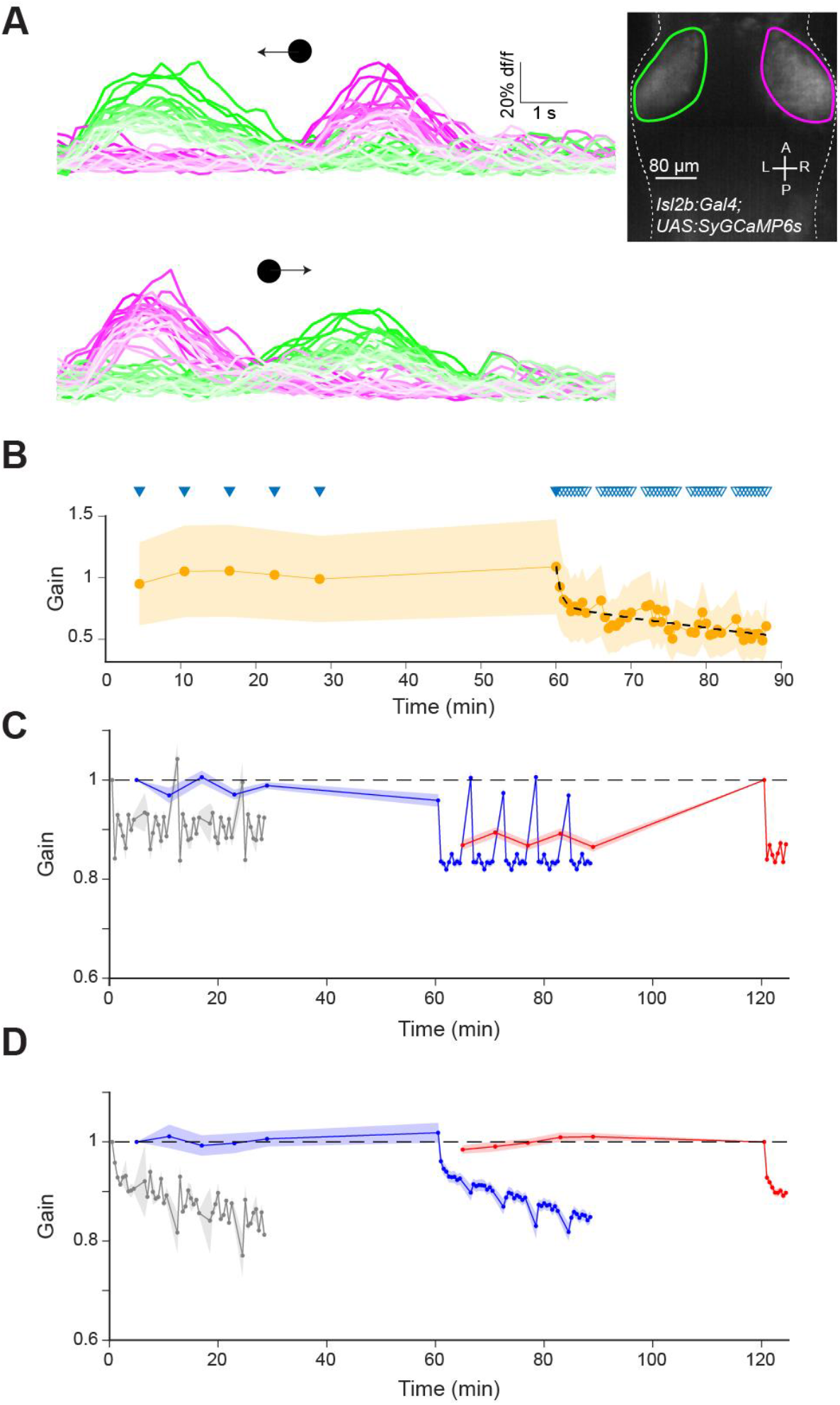
Imaging of retinal ganglion cell (RGC) axonal arbors and additional network simulations excluding retinal adaptation. **(A)** Calcium imaging of RGC axon terminals in OT in an *isl3:GAL4;UAS:SyGC6s* transgenic larva during leftward (top) and rightward (bottom) moving spot stimuli. Faint lines indicate later presentations. Inset: Mean z-projection of an image volume showing the left (green) and right (magenta) ROIs containing RGC axon terminals in the tectum. (**B**) RGC axon terminal responses (n=4 fish) normalised by the mean of the first two responses in each fish. Triangles indicate stimulus presentation times. Data from left and right OT were averaged. (**C**) Model prediction as per Figure 6, but without implementing RGC adaptation. (**D**) The net effect of simulated RGC adaptation, computed by taking the difference between the model prediction with and without RGC adaptation. Shaded areas indicate SEM.

