## Supplementary figures and images for "A recurrent network architecture explains tectal activity dynamics and experience-dependent behaviour"

### Supplementary File 1

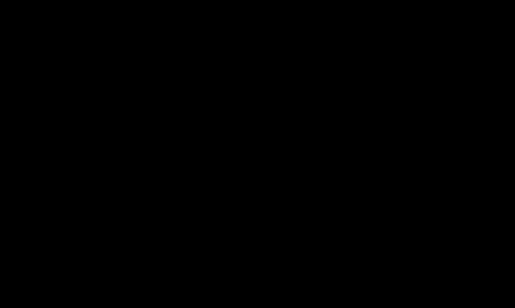
